# Transcranial random noise stimulation over the right prefrontal cortex does not improve performance on trained or untrained complex cognitive tasks

**DOI:** 10.64898/2026.04.10.717626

**Authors:** Scannella Sébastien, Riedinger Florine, Chenot Quentin

## Abstract

The present study aimed at evaluating the impact of high-definition transcranial random noise stimulation (HD-tRNS) applied to the right dorsolateral prefrontal cortex (DLPFC) on direct learning in computer-based complex tasks, and potential far transfer effects to a flight simulator task. Thirty young pilots in general aviation participated in a double-blind 11-week protocol that included a two-hour baseline session (week 1), 10 one-hour training sessions (weeks 2 to 6), a short-term (week 7) and a long-term (week 11) evaluations. Both stimulated, and sham groups exhibited improvements in trained (MATB and Space Fortress video game) and untrained (Flight Simulator) tasks from baseline to the first and last evaluation sessions. No significant differences between groups have been found either in terms of direct (trained tasks) or transfer (flight simulator and associated mental workload) effects. These findings contribute to the ongoing debate on the efficacy of transcranial brain stimulation for enhancing learning in healthy participants. Specifically, the present study demonstrates that the applied stimulation protocol yields no measurable benefit to learning processes, underscoring the need to explore alternative stimulation parameters and methodological approaches.

## Introduction

Enhancing cognitive performance—whether to gain a competitive edge, recover from injury, or adapt to increasingly complex operational demands—has long been a priority. Cognitive training has emerged as an effective approach to optimize brain function for complex tasks, with proven direct benefits [1–3]. Flying an aircraft for instance, is a widely considered complex task, especially when other factors such as failures or conflicts arise [4, 5]. In an attempt to mitigate errors and reduce cognitive workload, pilots are rigorously trained to cope with complex situations [6, 7]. Although these efforts have made aviation transportation the safest way to travel [8], there are still up to a thousand deaths per year in the last decade in commercial [9] and general aviation [10] due to issues in the management of critical situations. To some extent, this is also true in other complex systems, such as operating rooms, nuclear power plants, and more recently within the automotive domain with the emergence of autonomous vehicles [11]. As a consequence, understanding these phenomena and finding solutions to mitigate risks is important not only for aviation safety but also for all complex environments where lives are at stake.

There are now several practical approaches to help operators of complex systems stay in the loop and deal with critical situations. Among them, significant work has been done to improve training; more specifically, cognitive training. This research and development branch consists in working up front to improve brain resistance to external interferences that may have a negative impact on cognitive efficiency. To do so pilots are trained to cope with singular situations, which allows them to test their ability to maintain flight integrity [12]. These trainings however, have numerous limitations such as the fact that they cannot prepare to cope with unknown situations. As stated in several investigations, the occurrence of an incident or an accident is often the result of a unique combination of several unfavorable factors [13], leading to a novel and untrained situation. As a consequence, a complementary approach consists in training the brain in a more generic way to involve relevant cognitive functions more efficiently in a larger number of complex situations – including the ones that have never been encountered. It is important to distinguish between direct transfer, near transfer, and far transfer in cognitive training [14]. Direct transfer refers to improvements observed in the exact task that was trained. Near transfer occurs when training benefits a closely related task. For instance, practicing a standard N-back task may enhance performance on a dual N-back task, because they share common stimuli and they both rely on the same cognitive function: working memory. Far transfer involves improvements in more general or distinct tasks. For example, training in a dual N-back task may enhance performances in aircraft piloting. Like for the near transfer, the latter is based on the hypothesis that a far transfer effect will occur between the trained task and the target task due to shared cognitive processes [14].

With this in mind, researchers have developed several tasks that aim to train the brain generically to expect far transfer learning effects on real-life performance. Among them, executive functions, such as inhibition, memory updating, and cognitive flexibility [15] have often been targeted because they are highly recruited when dealing with complex and new situations [16, 17]. The Multi-Attribute Task Battery (MATB), for example [18], has been developed by NASA in an attempt to create such cognitive training support that would improve multitasking. In the same way, the Space Fortress (SF) video game [3, 19, 20] has been first developed to study learning in complex tasks [19] and later to train executive functions more dynamically with an additional level of entertainment [3]. SF was already used in 1994 as a mean to try to improve real flight performance in military pilots [21].

The promises of such transfer have led to numerous dedicated studies in healthy population [22, 23] and clinical domain [24, 25]. The benefits of such near and far transfer effects however, are not as high as expected. In a second-order meta analysis (*i.e*., a meta-analysis of meta-analyses), Sala and colleagues [14] revealed that once publication biases and placebo effects are accounted for, the presumed far-transfer benefits of cognitive training disappear: the inconsistencies between meta-analyses, showing none or marginal transfer effects resolve, and that the true variation among adjusted overall effect sizes vanishes. As a consequence, it seems that cognitive trainings alone are not powerful enough to expect far transfer effects – or even to induce large near transfer effects.

In an attempt to improve cognitive training and far transfer effects, a complementary approach emerged in the early 2000s by using non-invasive brain stimulation (NIBS) [26]. Among the different techniques, transcranial brain stimulation could be achieved using an electric field (Transcranial Electrical Stimulation; TES). In TES, two main approaches are distinguished: transcranial direct current stimulation (tDCS), which uses a constant current, and transcranial alternating current stimulation (tACS), which targets specific neuronal frequencies to modulate brain activity [27]. A third method that belongs to tACS is called transcranial random noise stimulation (tRNS). It employs a Gaussian-distributed variable-frequency alternating current to further enhance neuronal plasticity (see the review of Palaus and colleagues [28] for a description of all methods). The goal of tRNS is to target specific brain activity frequencies and to expect a modification in the discharge threshold of neurons in the related brain networks (*i.e*., a decrease of it). This, in turn, should induce a modification of behavioral performance [29–31]. Several studies have shown that it is possible to improve behavioral performance associated with high-level cognitive functions such as attention [32], working memory [33], inhibition [34], and multitasking [35]. In most of these studies, the dorsolateral prefrontal cortex (DLPFC) has been chosen as the stimulation target, due to his key role within the executive fronto-parietal network [36]. As with far transfer studies however, the NIBS literature remains divided on the actual benefits and effect sizes [37, 38].

To fully capture the effects of cognitive training, it is essential to move beyond behavioral performance alone and adopt a multimodal approach that integrates both subjective and objective measures of mental workload. Subjective assessment tools, such as the NASA-TLX [39], capture participants’ perceived effort, frustration, and task demand, providing valuable insights into the experiential impact of training. However, these self-reports are increasingly complemented by objective measures—such as performance metrics to offer a more robust evaluation of cognitive load. A particularly effective objective method involves using a secondary task, such as a concurrent auditory oddball paradigm during complex primary tasks (*e.g*., flight simulation [40]).

In this approach, participants perform an auditory oddball task (detecting infrequent target tones) while engaged in the primary task. Performance on the oddball task serves as an indicator of cognitive workload: degraded performance suggests high workload, as fewer attentional resources remain available for the secondary task. Recent studies have demonstrated that this method reliably reflects changes in cognitive load across different flight phases (e.g., cruise vs. landing), with performance on the oddball task significantly declining during high-demand phases compared to low-demand phases [41, 42].

In a previous study [43], we showed that the combination of a focal stimulation (*i.e*., HD-tRNS; 4 *×* 1 electrode montage) of the right DLPFC led to a better retention effect of skills acquired in the SF video game training compared to a placebo or a bilateral DLPFC stimulation. From these results, we designed the current hypothesis-driven study using the same HD-tRNS montage on the right DLPFC. We first wanted to reproduce our previous training results in SF with a longer training period. In addition, we wanted to evaluate a similar possible benefit of stimulation in the widely used MATB [18]. Our main goal was to assess possible transfer effects over a more ecological task: flying in a flight simulator. Both stimulated and sham groups were expected to perform better in the SF and MATB tasks after training (*i.e*., direct training effect).

This effect was expected to be associated with lower workload levels. However, our operational main hypothesis was that participants in the stimulated group would perform better (along with lower workload levels) in these trained tasks compared to those who received a placebo stimulation. Ultimately, flight simulator performance (*i.e*., far transfer effects) and associated workload levels, were also expected to be improved for the stimulated group compared to the sham one.

## Materials and methods

### Sample size estimation

The sample size estimation was based on our previous study [43], in which we observed a significant difference in SF performance between the sham and stimulated groups after one week of training (5 *×* 20 minutes of training; pre-post effect), with a moderate effect size (*η*² = .13; *Cohen’s f* = .387). As the present protocol extended the training duration to five weeks (10 *×* 60 minutes of training), we expected a larger between-group difference. Accordingly, we assumed a moderate-to-large effect size (*f* = .60) for the primary outcome (direct effect) at the end of the training. An *a priori* power analysis using *pwr.anova.test* (package ’*pwr* ’ [44]; *k* = 2 groups, *α* = .05, *f* = .60, *power* = .80) indicated a required sample size of 12 participants per group. To account for potential withdrawals, data quality issues, and to increase statistical power due to the measure of other variables (SF, MATB, Simulator performance), we planned to recruit 30 participants (15 per group).

### Participants

**Table 1.**
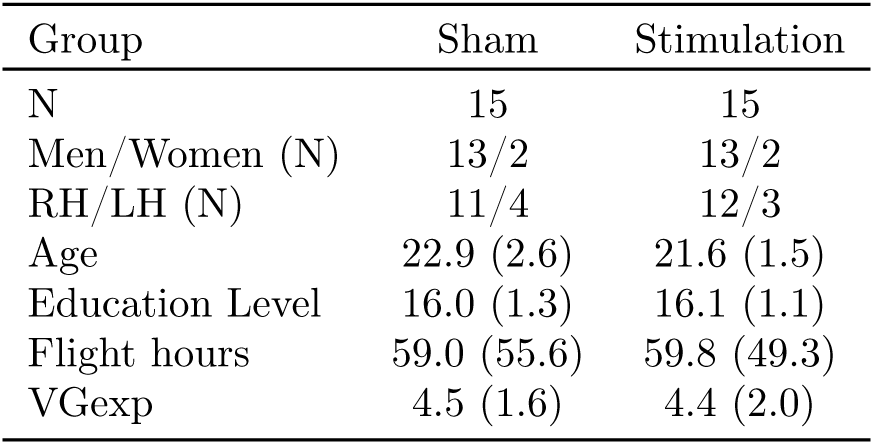
Demographic table. Mean (SD) of sample characteristics (unless marked “N”). RH/LH: Right/Left-Handed; VGexp: Video Game Experience Questionnaire

| Group | Sham | Stimulation |
| --- | --- | --- |
| N | 15 | 15 |
| Men/Women (N) | 13/2 | 13/2 |
| RH/LH (N) | 11/4 | 12/3 |
| Age | 22.9 (2.6) | 21.6 (1.5) |
| Education Level | 16.0 (1.3) | 16.1 (1.1) |
| Flight hours | 59.0 (55.6) | 59.8 (49.3) |
| VGexp | 4.5 (1.6) | 4.4 (2.0) |

31 young pilots in general aviation (Private Pilot License; mean flight hours=59.4; *SD* = 51.6) were recruited for this experiment in the vicinity of the ISAE-SUPAERO campus, Toulouse, France, through the means of flyers, social and professional networks and mailing-lists. The recruitment period lasted 17 months from April, the 1*^st^* 2022, to September, the 10*^th^* 2023. The experiment was approved by the local ethics committee (Montpellier University, IRB-EM: 2203C) and all participants signed informed consent prior to starting the experiment. One participant withdrew from the study after the first session due to concerns about potential adverse health effects of the stimulation. The remaining 30 participants included 26 men and 4 women; seven left-handed (mean age: 22.4; *SD* = 2.1; mean education level: 16 years; *SD* = 1.2). Participants received a €250 coupon once all sessions were completed.

#### Inclusion criteria and exclusion criteria

Inclusion and exclusion criteria were established prior to recruitment. Inclusion criteria were: age (18-35 yo); affiliation to social insurance; having read the information document about the experiment and signed the informed consent form; currently in training for, or already holding, a private pilot license (PPL). Exclusion criteria were: addiction (alcohol, drugs); major hearing loss; major visual deficit; including hemianopsia and color blindness; neurological or psychiatric pathology; known brain injury, drugs intake targeting the central nervous system.

### Experimental protocol

This study used a true experimental and longitudinal design with an intervention group and a control group, with outcomes being measured at multiple time points.

Participants were randomly assigned to conditions. The study employed a mixed design: group (tRNS stimulation vs. placebo) was a between-participant factor, and time was a within-participant factor, as participants were measured repeatedly across multiple time points. Data were collected in person in the laboratory for all sessions.

The 30 pilots participated in an 11-week protocol that included a two-hour baseline session (week 1), 10 one-hour training sessions (weeks 2 to 6), a one-hour short-term evaluation session (week 7), and a one-hour long-term evaluation session (week 11). The baseline, short- and long-term sessions were identical. In these sessions, the participants underwent 30 min of MATB, 30 min of SF and 1 hour of flight in the simulator. The flight scenarios were identical in terms of event timing and geographic environments through the three sessions. All other events have been strictly controlled to be equivalent across sessions. The ten training sessions were all identical. Twice a week, keeping the day and the time of the session as constant as possible, the pilots were trained in MATB and SF tasks together with a non-invasive brain stimulation (real or placebo) of the right DLPFC (see Figure 1).

**Fig 1.**
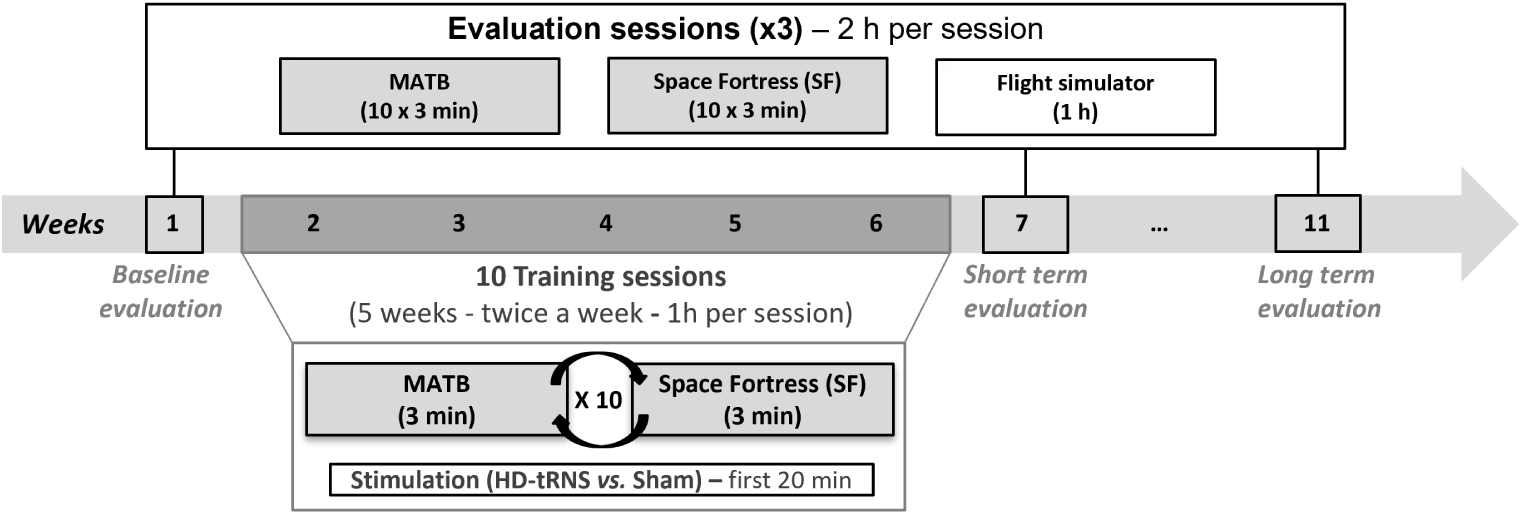
Study timeline. Participants enrolled for 11 weeks in the following 13 sessions. Session 1 (week 1): baseline for performance assessment in MATB, SF and Flight simulator. Sessions 2 to 11 (weeks 2 to 6): 10 training sessions in MATB and SF with real HD-tRNS on rDLPFC or placebo stimulation. Sessions 12 and 13 (week 7 and 11): short- and long-term performance evaluations.

Baseline Flight simulator and workload scores were published in a previous article [45] to develop a flight simulator score. All other data including, performance in trained tasks, the training and longitudinal results reported here are new.

### Tasks

#### Trained tasks: MATB-II and Space Fortress

For the ten training sessions (sessions 2 to 11), two tasks were used, the MATB and Space Fortress video game.

*The MATB-II* [18] is a computer-based task in which participants must handle different tasks (see Figure 2.a). We used four subtasks: (1) System monitoring (reporting deviant values in six different scales), (2) Tracking (keep the moving crosshair in a central area using a joystick), (3) Radio communications (set radio frequency values according to auditory messages) and (4) Resource management (manage fuel pump status to optimize tank levels). For this study, a Matlab-modified version of the NASA MATB-II (https://github.com/VrdrKv/MATB) was used. A regular AZERTY keyboard and a Logitech Extreme 3D Pro joystick were used as controllers. Performance on this task was evaluated with a global score averaging all sub-tasks (see).

**Fig 2.**
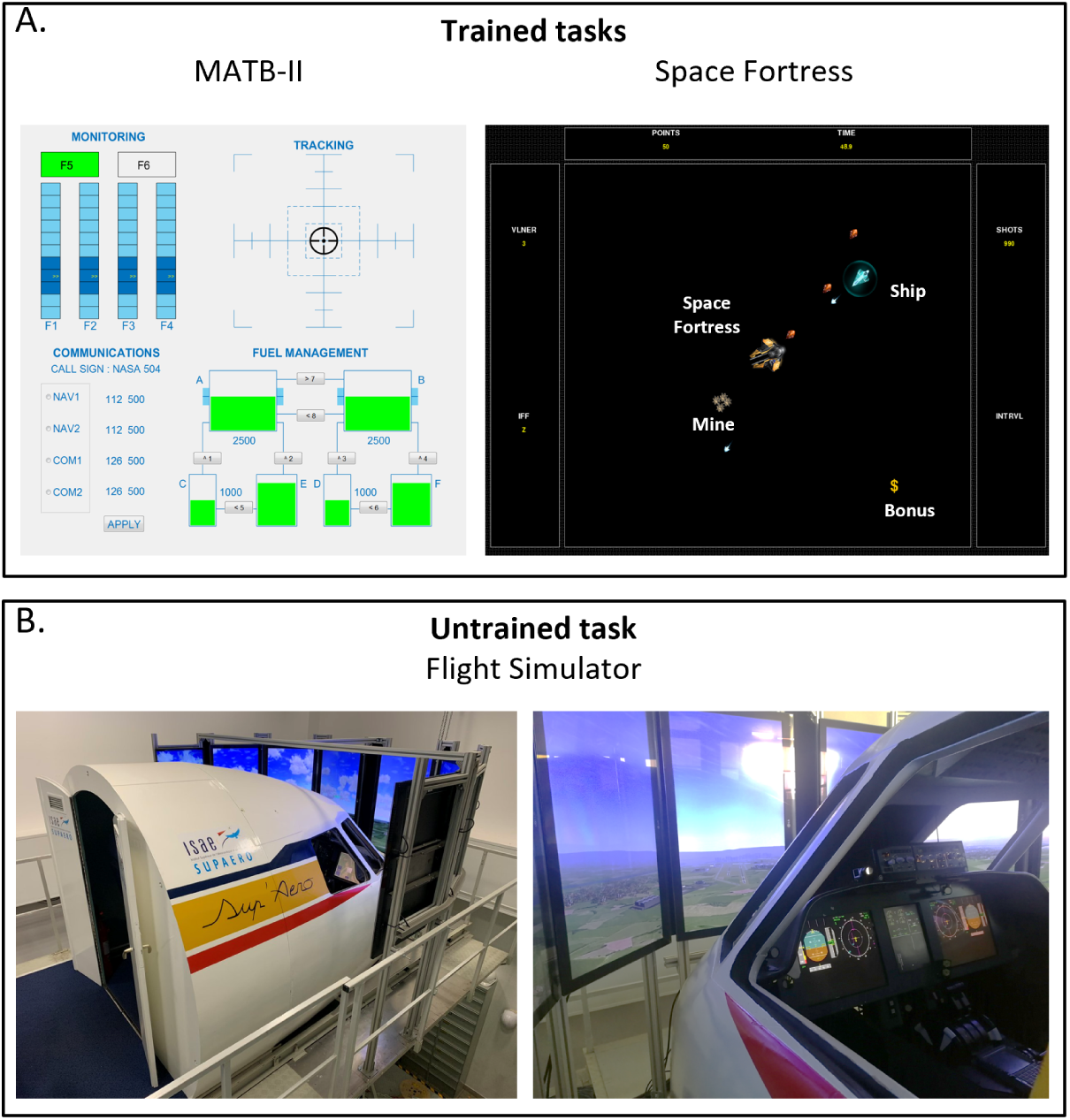
A. Trained tasks (left: MATB-II; right: Space Fortress). B. Untrained task: 3-axis flight simulator “pegase” of ISAE-SUPAERO.

*Space Fortress* [19] is an arcade-style 2D game that can be decomposed into four sub-tasks: (1) Controlling your spaceship, (2) Destroying the space fortress (which tries to destroy your ship), (3) Identifying and destroy two types of incoming mines, and (4) Capturing bonuses (identify a predefined two-symbol sequence among others). It is considered a multitasking game (see Figure 2.a) since the participants must operate all subtasks simultaneously to perform well (i.e., to get the highest score). A complete description of this task and rules is available in the work of Shebilske and colleagues [46]. For the current experiment, we used the SF version 5.1.0 running with Python 2.7 (https://github.com/CogWorks/) and commands were managed with a regular AZERTY keyboard. Performance on this task was evaluated with a global score (see section).

#### Untrained task: Flight simulator

During the flight simulator sessions (baseline, short- and long-term sessions), pilots were installed in the ISAE-SUPAERO flight simulator. This simulator is a 3-axis flight simulator (*Pegase*, ISAE-SUPAERO, see Figure 2B). Eight screens arranged in a 180° arc displayed the environment outside the cockpit. In the cockpit, five screens displayed flight parameters, with a display similar to that of an A320 Airbus. The cockpit included Airbus-like stick and throttle (two engines), and a Flight Control Unit (FCU) panel (deactivated for this experiment). The simulation software was FlightGear 2.4, and the simulated aircraft was an A320. The aircraft’s flight-by-wire controls and aerodynamics were homemade, based on the A320’s systems and flight envelope.

Simulator data were recorded at 50 Hz including time (seconds), geographic coordinates (latitude and longitude in decimal degrees), speed (knots), altitude (feet), variometer (feet per minute), g-force and aircraft axes (pitch, roll, yaw in degrees). Four flight scenarios were created according to flight rules (Visual Flight Rules; VFR *vs.* Instrument Flight Rules; IFR) and difficulty (Low *vs.* High). The four scenarios consisted of flight traffic patterns in the vicinity of Toulouse-Blagnac airport (France), and the difficulty was manipulated according to visibility, landing type and failure (see Table 2, Figure 3 and [45] for more details). Pilots’ performance was assessed with a composite score taking into account five criteria: time difference; target speed and altitude deviation, flight path deviation; distance to landing point and landing G-force (see).

**Fig 3.**
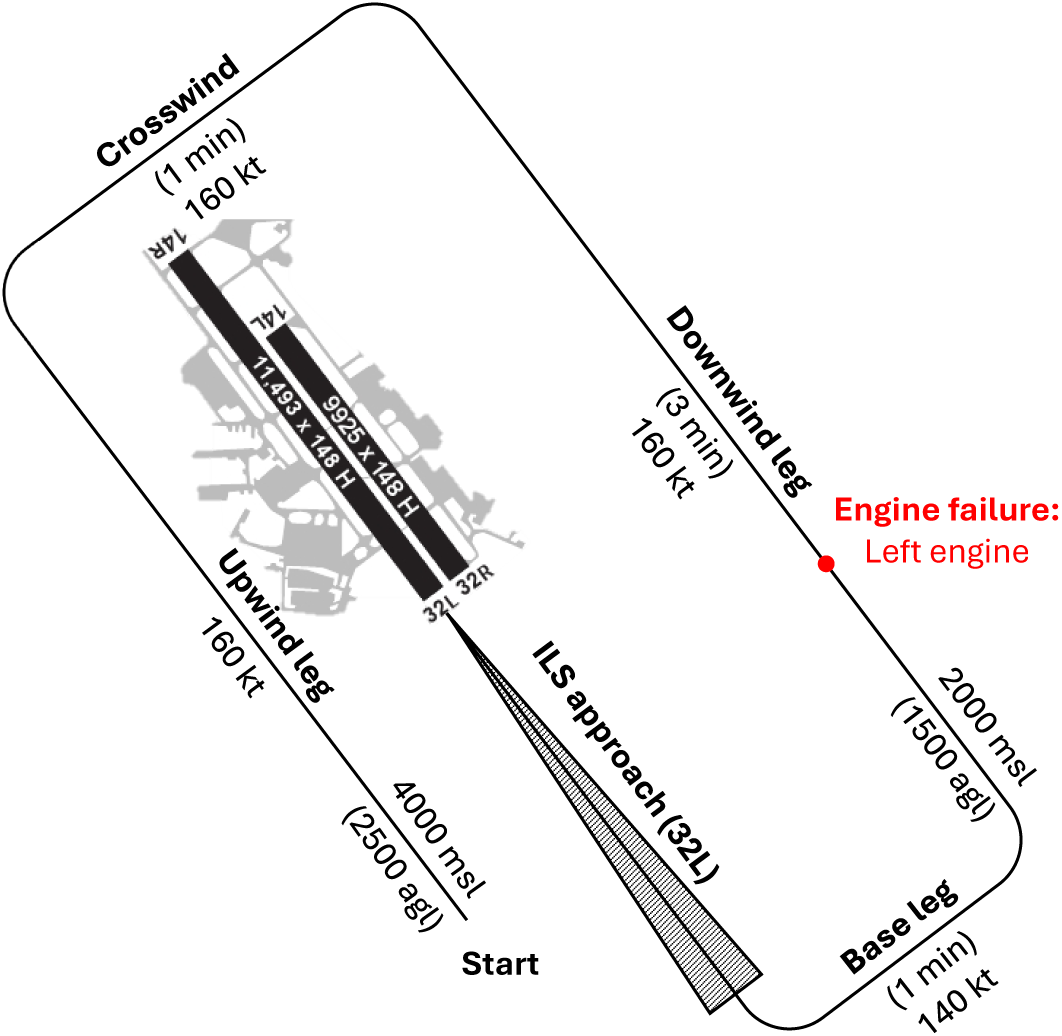
Illustration of a standard traffic pattern scenario (IFR-high) used across the four different flight scenarios (not to scale).

**Table 2.** Flight Simulator Scenarios. VFR: Visual Flight Rules. IFR: Instruments Flight Rules. ILS: Instrument Landing System. Alt: Altitude.

| Type | Difficulty | Visibility | Landing | Failure |
| --- | --- | --- | --- | --- |
| VFR | Low | Clear | Visual | None |
|  | High | Cloudy | Visual | Alt+Speed |
| IFR | Low | Clear | ILS | None |
|  | High | Fog+night | ILS | Engine |

#### Untrained task: Auditory oddball

During the flight simulator scenarios, participants concurrently performed an auditory oddball task. The task consisted of standard low-frequency tones (750 Hz pure tones, 78 dB SPL) and target high-frequency tones (1250 Hz pure tones, 78 dB SPL). Each tone had a duration of 100 ms and were presented in pseudo-randomized sequences with a fixed proportion of 20% target tones (2 targets per 10 stimuli). The inter-stimulus interval (ISI) had a mean duration of 1.25 s and included a temporal jitter of *±* 0.75 s, resulting in ISIs ranging from 0.5 s to 2.0 s. Given that flight scenarios lasted approximately 10 min, each scenario contained about 480 auditory stimuli, including 96 target tones. Participants were instructed that the piloting task was the primary task and to respond to as many target tones as possible by pressing the joystick main trigger.

### Transcranial random noise stimulation

The tRNS was applied only during the ten training sessions (weeks 2 to 6). Two groups were created and participants were randomly assigned to one of them at the beginning of the experiment. The study was conducted as a double-blind trial; therefore, both the participants and the experimenter were unaware of the stimulation condition. This was achieved using the password-protected double-blind procedure embedded in the Neuroelectrics NIC software (ver. 2.1.3.7). As a result, participants were assigned to group A or group B. The actual group was revealed after the statistical analyses were completed. Both groups were equipped with a neoprene cap with a 4 *×* 1 electrode montage, with 5 circular (*π* cm²) NG Pistim gel electrodes placed on C4, Fp2, Fz, F8 locations for the peripheral electrodes and F4, over the right DLPFC in the 10-20 international system, as the central electrode (see Figure 4). Note that because we were using an alternating current without DC-offset, the notion of active and return electrodes is not relevant here.

**Fig 4.**
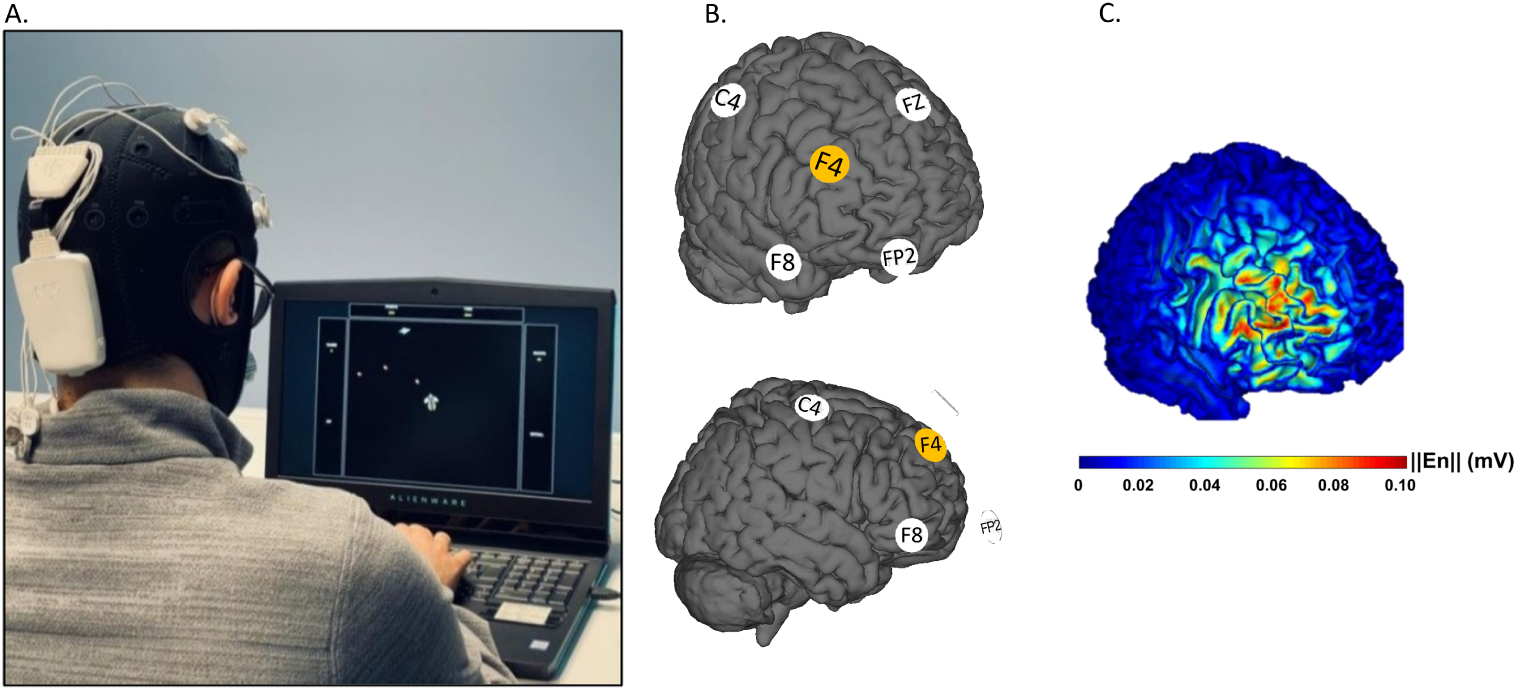
A. Participant training on Space Fortress while receiving tRNS. B. A 4 *×* 1 electrode montage has been used with central electrode F4 (orange) as the stimulation electrode over the right DLPFC and Fz, FP2, F8 and C4 electrodes as return electrodes (white). C. Current density distribution of the active simulation over the right DLPFC white matter (extracted from NIC v2.4 software; Neuroelectrics).

The tRNS waves for the stimulated group (HD-tRNS) were generated using a Starstim device (Neuroelectrics). The random level of current was generated for every sample (sampling rate 1280 samples/s). The current intensity was randomized between *−*0.5 and 0.5 mA (mean = 0; SD = 0.333) following a normal distribution, in line with previous studies [35, 47]. As the current distribution in a large band of high frequencies has been recommended for tRNS [48], we chose the largest one allowed by our device, distributed between 100 and 500 Hz. In the 4 *×* 1 HD-tRNS montage, the central electrode (F4) always sent or received 100% of the delivered current, while the 4 peripheral electrodes sent or received 23% for F8 and Fz and 27% for C4 and Fp2 of the current (to account for to the distance from the central electrode). Both groups received a gradual rise (current ramp up) of 15 seconds at the beginning of the stimulation and a gradual decrease (current ramp down) during the last 15 seconds. Between ramps, 20 min of stimulation matched the first 20 min of the training session. The stimulated group received a total of 394.6 mC during the whole stimulation, distributed mainly on the targeted right DLPFC. The sham group received no stimulation during the 20 min of SF and MATB training. Consequently, the sham group received a total of 7 mC including both ramps.

### Behavioral Measures

#### Questionnaire

##### Demographic questionnaire

Participants were asked to report their sex, age, native language, handedness and education level.

##### Video game experience questionnaire

Participants provided information about current and past video game habits. A composite score (VGexp) representing their global level from 1 to 10 was calculated from three subscores: number of game sessions per year, number of game hours per day, and whether they consider themselves playing it intensively.

##### Flight experience questionnaire

Participants reported their total flight hours in real aircraft as well as their flight hours in flight simulators.

##### tRNS Adverse Effects Questionnaire

A translated version (French) of the tDCS Adverse Effects Questionnaire [49] was administered to the participants (renamed tRNS Adverse Effects Questionnaire in this article) before and after each of the training session. This questionnaire included 12 items on how they currently evaluate different sensations (headache, difficulty to concentrate, change of mood, change in visual perception, tingling, itching, burning, pain, fatigue, nervousness, unpleasant sensations and nausea) in which the participants rated on a five-point Likert scale (from 0 to 4). The total score of this questionnaire was computed as the sum of all responses and ranged from 0 to 48.

### Trained tasks

#### Space Fortress Score

An SF score was computed for each run, with points allocated as in Table 3. To facilitate interpretation and comparability, the scores of all participants and runs were transformed into a z-score. The data followed a normal distribution.

**Table 3.** Space Fortress score distribution.

| Subtask | Rule | Points |
| --- | --- | --- |
| Flying | - Being hit | -35 |
|  | - Being destroyed (3 hits) | -100 |
|  | - Crossing game borders | -35 |
|  | - Colliding with the Space Fortress | -35 |
| Fortress | - Destroying the Space Fortress | 250 |
| Mine | - Destroying a Type-1 mine | 50 |
|  | - Destroying a Type-2 mine (which must first be identified) | 60 |
|  | - Failing to destroy a mine before it disappears | -50 |
| Bonus | - Capturing a missile bonus | 50 |
|  | - Capturing a point bonus | 100 |

#### MATB Score

A composite MATB score was calculated for each run by aggregating performance across the three subtasks (RESMAN, TRACKING and RESMAN). For resource management, performance was defined as the difference between the target value (*e.g.*, 2500) and the participant’s mean deviation from that target. Tracking performance was quantified as the participant’s mean distance from the center of the tracking display. Monitoring performance was defined as the mean response time to alarm events. All three measures were log-transformed to reach normality and z-scored across all runs and participants. The overall MATB score was then computed as the mean of these three z-scores.

### Untrained tasks

#### Flight Simulator Score

The Flight Simulator Score also consisted of an aggregate of the performance in four different flight scenarios (see the work of Chenot et al., [45]).

For each scenario, five metrics were calculated:

1. Time Metric: Defined as the absolute difference in seconds between the expected and actual time taken for each flight phase. For instance, the time metric for the downwind phase was calculated based on a reference duration of 3 min.
2. Speed Metric: Similar to the altitude metric, this was determined by the absolute area under the curve, comparing the expected (e.g., 140 knots) and achieved speeds.
3. Altitude Metric: Computed as the absolute area under the curve, representing the deviation from the expected altitude (e.g., 2000 feet) to the altitude flown by the pilot.
4. Approach Metric: Calculated as the mean distance deviation between the expected approach path (as it would be executed by an autopilot) and the pilot’s actual flight path.
5. Landing Metric: This metric involved two components: the measured g-forces during landing and the distance deviation from the ideal landing spot (as determined by an automated pilot landing) to the pilot’s landing position.

In order to compute the score for each scenario, a three-step preprocessing of these metrics was performed using R studio (ver. 4.2.2 [50]), allowing us to average all five metrics.

1. Log transformation: Applied to address the positive skewness observed in the data distribution of these metrics;
2. Normalization (Z-Score transformation): Data were then normalized and converted into z-scores.
3. Inversion: these scores were then inverted, ensuring that positive scores corresponded to superior performance. For example, the pilot with the highest area under the curve (*i. e.* largest deviation) for the altitude metric had its positive z-score inverted into a negative z-score.

Given the specificities of the four traffic patterns (target altitude, speed, etc.), the data preprocessing pipeline (log-transformation and z-scoring) was applied to each scenario independently, but taking into account the three sessions (baseline, short and long-term). This approach ensured that an individual score was computed for each scenario and each session. Finally, the Flight Simulator Score for each session was calculated by averaging the z-scored performance scores of all scenarios completed within that session.

#### Workload Score

Participants’ workload was assessed using both subjective and objective measures. Subjective workload was evaluated with the NASA-TLX questionnaire [39], administered after each of the four flight scenarios during the evaluation sessions (*i.e.*, baseline, short-term, and long-term). Objective workload was measured by the percentage of missed auditory target tones in the concurrent oddball task performed during the flight scenarios. This metric provided an estimate of the remaining available auditory attentional resources in each scenario.

### Hypotheses and statistical analyses

Three hypotheses were formulated to assess the effects of tRNS on learning and performance across the trained (SF, MATB) and untrained tasks (flight simulator).

#### Hypothesis 1: Direct effects of tRNS on trained task learning rates

We first hypothesized that the tRNS group would exhibit better learning rates in trained tasks than the SHAM group. This hypothesis was tested separately for the SF (H1a) and MATB (H1b) tasks.

Operationally, training sessions only (T01–T10) were used to compute individual learning rates. For each participant, performance scores were regressed onto the natural logarithm (ln) of session number, and the coefficient of the ln(Session) term was taken as an index of learning rate. Group differences in learning rate were assessed using independent-samples *t* -tests for each task.

#### Hypothesis 2: Direct effects of tRNS on post-training trained task performance

The second hypothesis postulated that the tRNS group would exhibit higher training effects compared to the SHAM group on the trained tasks: (H2a) *Space Fortress* and (H2b) *MATB*. Performance was analyzed in the three evaluation sessions: baseline, short-term, and long-term. For each outcome, we fitted a linear mixed-effects model of the form of equation 1.

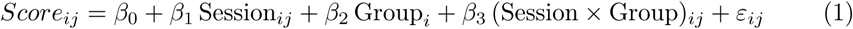

where *ε_ij_*is the residual error.

#### Hypothesis 3: Far transfer effects of brain stimulation on untrained tasks

The third hypothesis postulated that the tRNS group would exhibit better performance in the untrained task (flight simulator score, H3a); and lower subjective (H3b) and objective (H3c) workloads. For each variable, we applied linear mixed-effects models using the same specification described in equation 1 to evaluate the effects of Session (Baseline *vs.* Short-term *vs.* Long-term) and Group (real tRNS *vs.* Placebo).

#### Data processing and statistical analyses

Data processing, including data transformation, learning rates calculation, statistical analyses and plot generation were made with Python v3.13.5 and R version 4.3.1 in Visual Studio Code v1.111.0.

For the five linear mixed models, assumptions were evaluated for autocorrelation (Durbin-Watson, DW), heteroscedasticity (Breusch-Pagan, BP), and normality (Shapiro-wilk, SW) on model residuals. The SF model met residual assumptions (DW = 2.053, BP *p* = .225, SW *p* = .679). The NASA-TLX model met residual assumptions (DW = 2.052, BP *p* = .818, SW *p* = .208) as well as for the Oddball model (DW = 2.369, BP *p* = .153, SW *p* = .737). The Flight Simulator model showed acceptable residual normality (SW *p* = .603) and no autocorrelation concern (DW = 2.483), but a possible heteroscedasticity issue (BP *p* = .036). The MATB model showed residual assumption deviations, including non-normality (SW *p* = .017) and heteroscedasticity (BP *p* = .0037), with no autocorrelation concern (DW = 2.151). To account for heteroscedasticity of residuals, robust standard errors were computed post hoc using the sandwich estimator CR2 [51] applied to the linear mixed model, as implemented in the *clubSandwich* R package.

## Results

### tRNS adverse effect questionnaire

The mean tRNS adverse effect questionnaire score before the session was 1.61 (SD = 1.19) for sham group and 1.96 (SD = 1.62) for stimulation group. After the session, mean scores increased to 2.30 (SD = 1.41) for the sham group and 3.06 (SD = 1.90) for the stimulation group. A repeated-measures ANOVA examining the score with Pre-Post session as a within-subject factor and Group as a between-subject factor revealed a significant main effect of Pre-Post session (*F* (1, 29) = 22.40, *p < .*001, *η*^2^ = .44), indicating an increase in scores after the session. The main effect of Group was not significant (*F* (1, 29) = 1.28, *p* = .27, *η*^2^ = .04), suggesting no overall difference between groups. Importantly, the Group *×* Pre-Post session interaction was not significant (*F* (1, 29) = 1.20, *p* = .28, *η*^2^ = .04).

### Trained tasks

#### Assessment of potential covariates

No significant correlation between flight hours and initial or final performance in the trained tasks has been found (best *R*^2^ = .06; *p* = .14). Video game experience, however, was significantly correlated to initial performance in MATB (*R*^2^ = .18; *p < .*05) and SF (*R*^2^ = .23; *p < .*01) as well as to long term performance (i.e., 1 month after training) in MATB (*R*^2^ = .16; *p < .*05) and SF (*R*^2^ = .29; *p < .*01). As a consequence, the VGexp has been kept as a covariate factor in subsequent analyses.

#### Hypothesis 1: Effect of tRNS on learning rates of trained tasks

No significant differences between groups have been found in the learning rates during the MATB training (*t* = 1.29; *p* = .21; *mean* = 0.363; *SD* = 0.174 for the tRNS group and *mean* = 0.303; *SD* = 0.131 for the sham group; see Figure 5.A). Similarly, non-significant results were obtained for the SF learning rate (*t* = *−*0.65; *p* = .52; *mean* = 0.713; *SD* = 0.245 for the tRNS group and *mean* = 0.757; *SD* = 0.230 for the sham group; see Figure 5.B). These results indicate that, contrary to our hypothesis 1, the HD-tRNS applied over the right DLPFC did not improve learning rates during the training period. In addition, the VGexp was not a significant predictor of the learning rates for both MATB (*t* = *−*0.04; *p* = .97) and SF (*t* = 1.96; *p* = .06).

**Fig 5.**
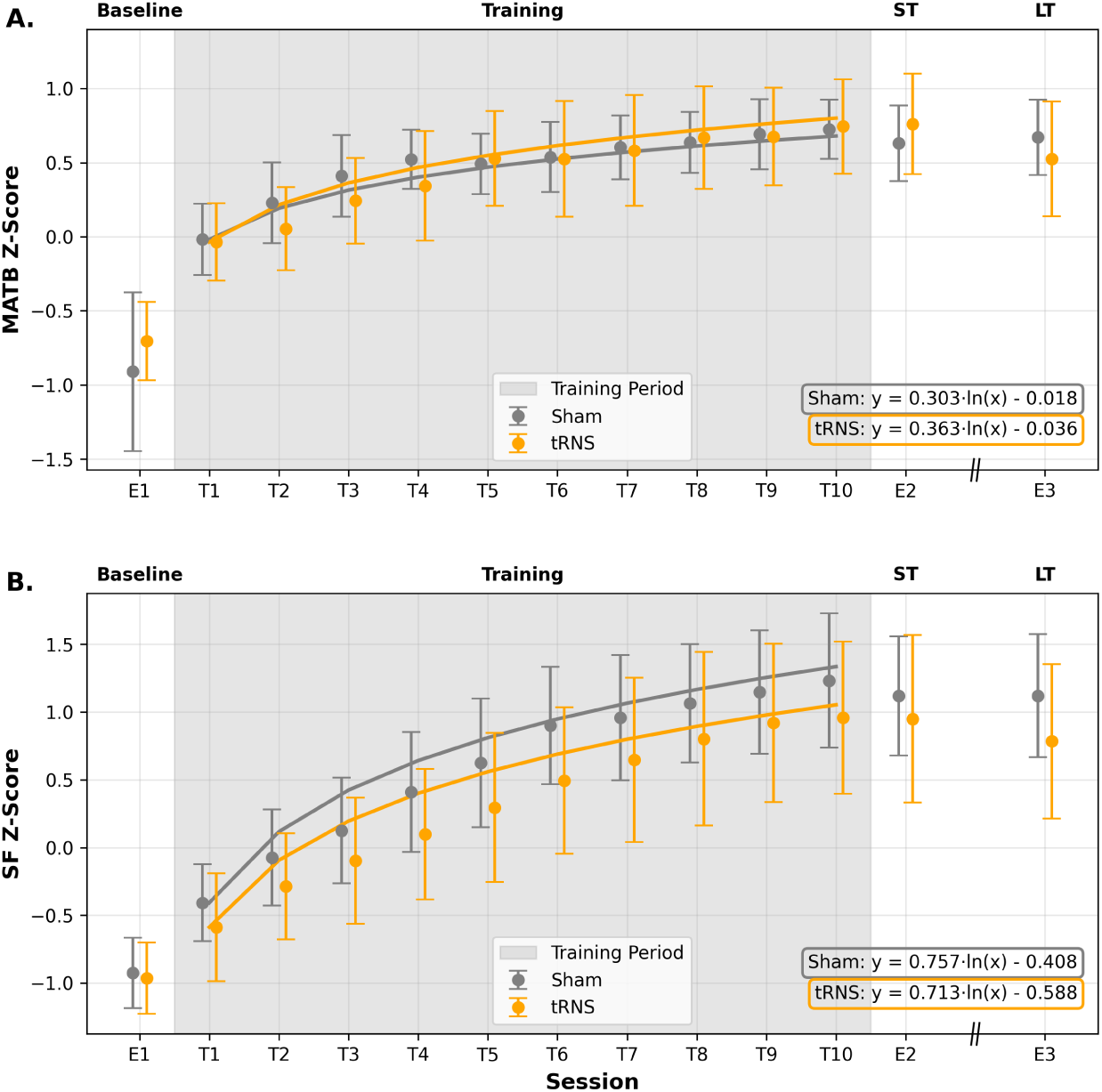
MATB (A) and Space Fortress (B) scores through the 13 sessions during 11 weeks and logarithmic fit during the training period. E1, E2, E3: Evaluation sessions; T1-T10: training sessions. ST: short term; LT: long term. Vertical bars represent the 95% confidence interval.

### Hypothesis 2: Effect of tRNS on post-training trained task performance

#### MATB

The linear mixed-effect model regression of MATB scores showed no significant group effect (*Z* = 0.80; *p* = .42; see Figure 6.A and Table 4). Both groups benefited from the training as attested by the significant Session effects between baseline and short-term evaluations (*Z* = 10.91; *p < .*001) and between baseline and long-term evaluations (*Z* = 11.20; *p < .*001). In addition, the video-game experience significantly explained the overall MATB scores (*Z* = 2.67; *p < .*01).

**Fig 6.**
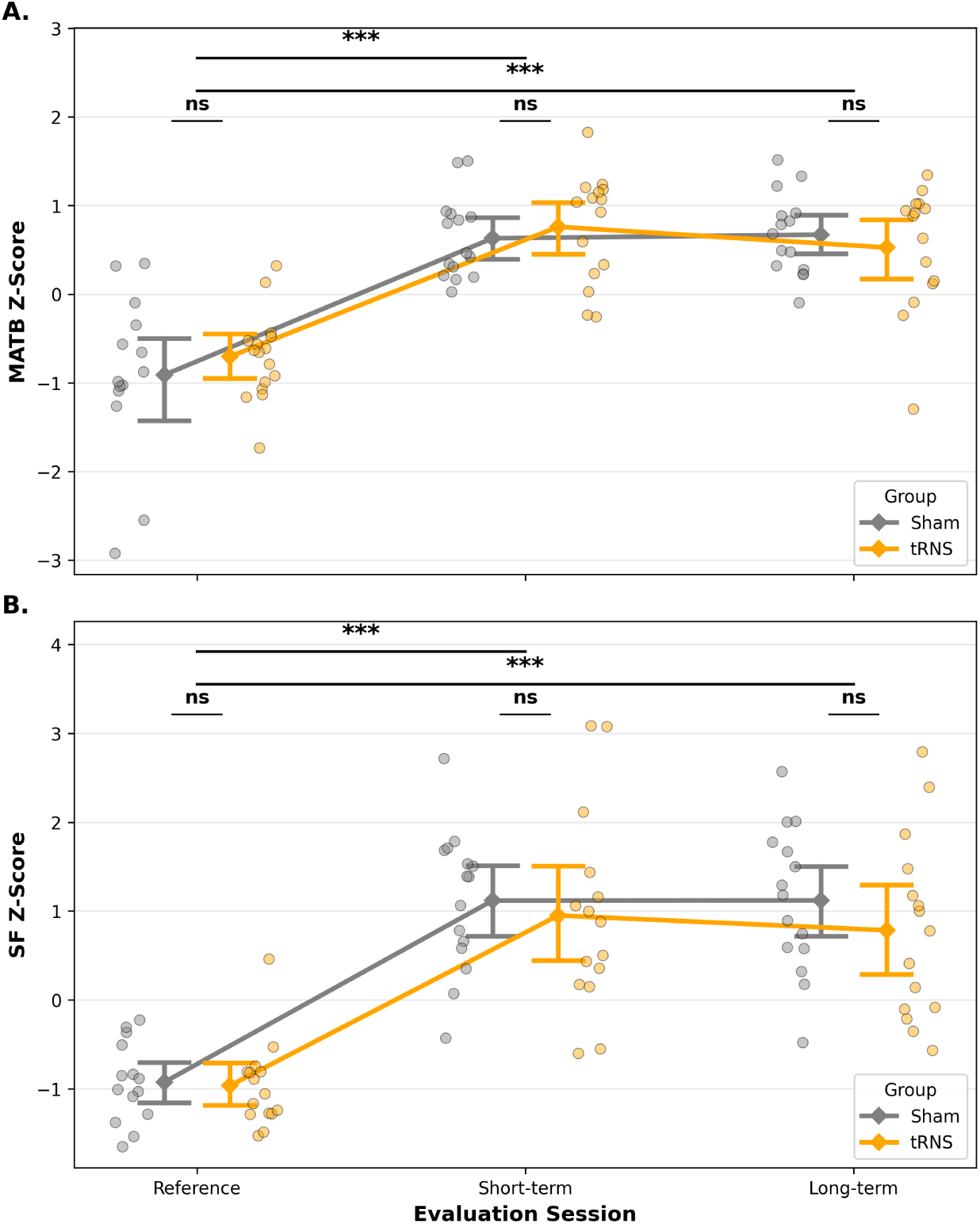
MATB (A) and SF (B) scores for the reference (E1), the short-term (E2) and the long-term (E3) sessions. Vertical bars represent the 95% confidence interval. ***: *p < .*001; ns: non-significant.

**Table 4.** Mixed-effects model results for MATB and SF performance with gaming experience (VGexp) as covariate. Reference level: baseline session (E1).

| Predictor | MATB Performance |  |  |  | SF Performance |  |  |  |
| --- | --- | --- | --- | --- | --- | --- | --- | --- |
| | $\beta$ | 95% CI | $z$ | $p$ | $\beta$ | 95% CI | $z$ | $p$ |
| Intercept | -1.54 | [-2.10, -0.98] | -5.42 | < .001 | -1.97 | [-2.69, -1.24] | -5.31 | < .001 |
| Session: short-term | 1.54 | [ 1.26, 1.81] | 10.91 | < .001 | 2.02 | [ 1.73, 2.32] | 13.40 | < .001 |
| Session: long-term | 1.58 | [ 1.30, 1.85] | 11.20 | < .001 | 2.03 | [ 1.73, 2.32] | 13.41 | < .001 |
| Group (tRNS vs. Sham) | 0.18 | [-0.25, 0.60] | 0.80 | .421 | -0.01 | [-0.55, 0.53] | -0.04 | .969 |
| Short-term × Group | 0.01 | [-0.39, 0.40] | 0.03 | .973 | -0.14 | [-0.56, 0.28] | -0.65 | .515 |
| Long-term × Group | -0.28 | [-0.67, 0.12] | -1.38 | .167 | -0.30 | [-0.72, 0.12] | -1.40 | .162 |
| VGexp | 0.14 | [ 0.04, 0.25] | 2.67 | .008 | 0.24 | [ 0.10, 0.37] | 3.36 | .001 |

95% CI: confidence interval. Bold $p$ : significant at $p < .05$ .

#### Space Fortress

Similarly, the linear mixed-effect model regression of SF scores showed no significant group effect (*Z* = *−*0.27; *p* = .97; see Figure 6.B and Table 4). Both groups benefited from the training as attested by the significant Session effects between baseline and short-term evaluations (*Z* = 13.40; *p < .*001) and between baseline and long-term evaluations (*Z* = 13.41; *p < .*001). In addition, the video-game experience also significantly explained the overall SF scores (*Z* = 3.36; *p < .*01).

### Untrained tasks

#### Assessment of potential covariates

No significant correlations were found between video game experience and NASA-TLX, Oddball, or simulator scores at either baseline or post-training sessions (best *R*^2^ = .05; *p* = .24). In contrast, prior light aircraft flight experience predicted post-training simulator performance, with greater flight hours associated with better outcomes (best *R*^2^ = .25; *p* = .005). No other significant associations were observed for NASA-TLX or Oddball scores. Accordingly, flight experience (light aircraft hours) was included as a covariate in subsequent analyses of simulator performance.

### Hypothesis 3: Effect of tRNS on post-training simulator performance and workload assessments

#### Flight Simulator Performance

The linear mixed-effect model regression of simulator performance showed no significant group effect (*Z* = *−*0.38; *p* = .7; see Figure 7.C and Table 5). Both groups demonstrated significant performance improvements, particularly at the long-term evaluation (*Z* = 3.48; *p < .*001) compared to baseline. As expected from the correlation analyses, flight experience significantly predicted simulator scores (*Z* = 2.43; *p < .*05), with experienced pilots showing better flight simulation scores.

**Fig 7.**
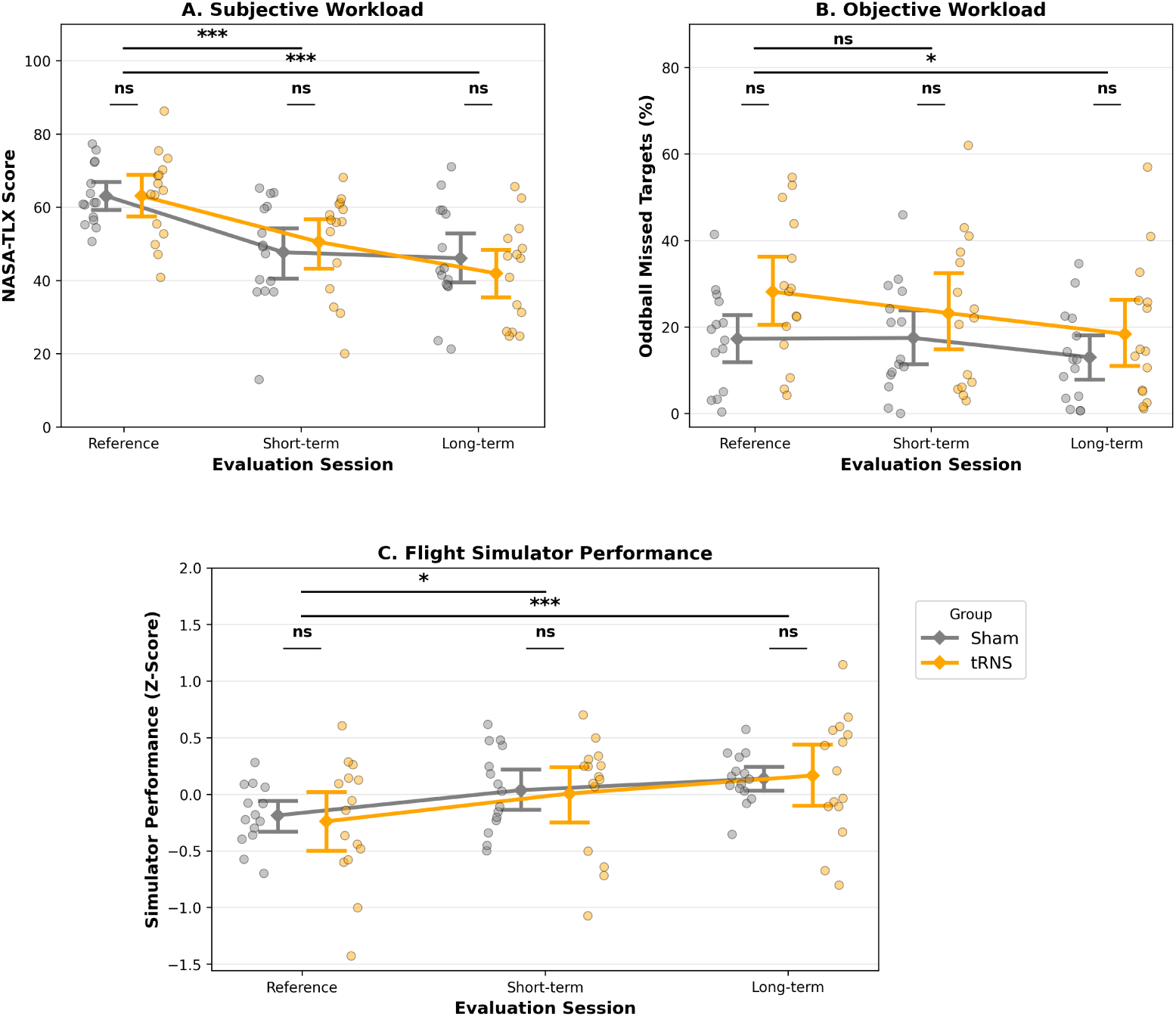
Untrained task scores for subjective (A) and objective (B) workloads and Flight Simulator performance (C). Vertical bars represent the 95% confidence interval. *: *p < .*05; ***: *p < .*001; ns: non-significant.

**Table 5.**
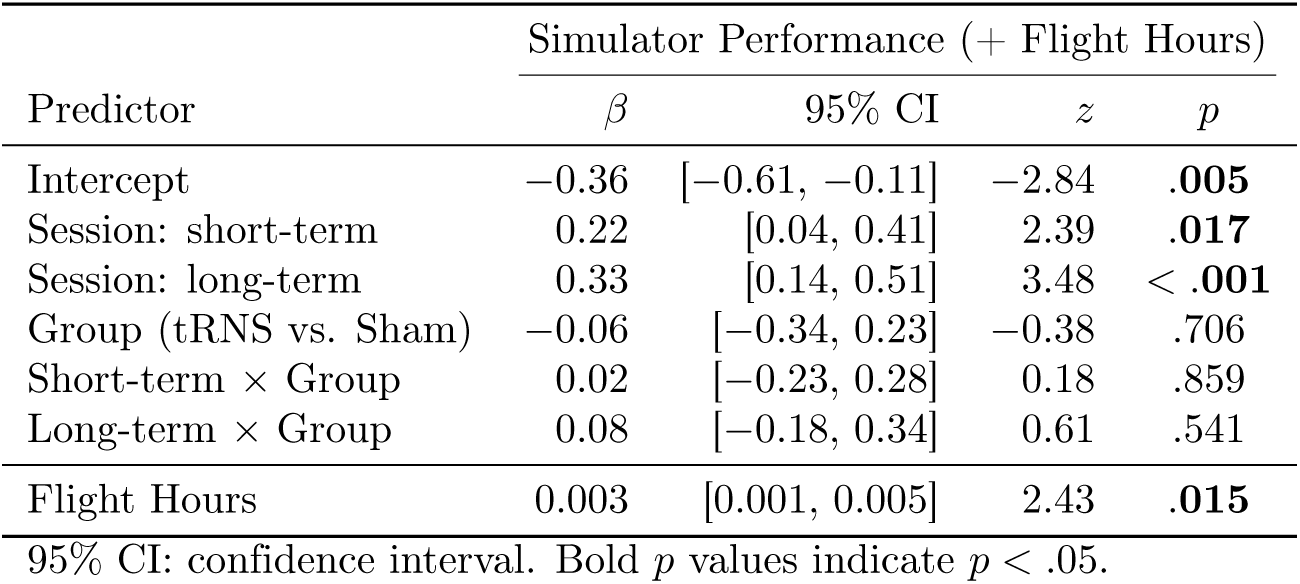
Mixed-effects model results for simulator performance (with flight hours as covariate). Reference level: baseline session (E1).

#### Subjective Workload (NASA-TLX)

Similarly, the linear mixed-effect model regression of NASA-TLX scores showed no significant group effect (*Z* = 0.01; *p* = .99; see Figure 7.A and Table 6). Both groups exhibited significant reductions in workload across sessions, for both the short-term (*Z* = *−*4.94; *p < .*001) and the long-term (*Z* = *−*5.46; *p < .*001) compared to baseline.

**Table 6.** Mixed-effects model results for subjective workload (NASA-TLX) and objective workload (Oddball missed targets). Reference level: baseline session (E1).

| Predictor | NASA-TLX |  |  |  | Oddball Missed Targets |  |  |  |
| --- | --- | --- | --- | --- | --- | --- | --- | --- |
| | $\beta$ | 95% CI | $z$ | $p$ | $\beta$ | 95% CI | $z$ | $p$ |
| Intercept | 63.06 | [56.52, 69.59] | 18.90 | < **.001** | 18.03 | [10.62, 25.44] | 4.77 | < **.001** |
| Session: short-term | -15.36 | [-21.46, -9.26] | -4.94 | < **.001** | -0.58 | [-4.84, 3.69] | -0.27 | .791 |
| Session: long-term | -17.00 | [-23.10, -10.90] | -5.46 | < **.001** | -5.02 | [-9.29, -0.76] | -2.31 | **.021** |
| Group (tRNS vs. Sham) | 0.03 | [-9.22, 9.27] | 0.01 | .995 | 10.14 | [-0.29, 20.58] | 1.91 | .057 |
| Short-term × Group | 2.79 | [-5.83, 11.42] | 0.63 | .526 | -4.40 | [-10.36, 1.55] | -1.45 | .147 |
| Long-term × Group | -4.14 | [-12.76, 4.49] | -0.94 | .347 | -4.79 | [-10.75, 1.16] | -1.58 | .115 |

95% CI: confidence interval. Bold $p$ values indicate $p < .05$ .

#### Objective Workload (Oddball Missed Targets)

Finally, the linear mixed-effect model regression of Oddball missed targets also showed no significant group effect (*Z* = 1.91; *p* = .06; see Figure 7.B and Table 6). Objective workload improved significantly across sessions, only at the long-term evaluation (*Z* = *−*2.31; *p < .*05). Flight experience (flight hours) was not a significant predictor of flight performance (*Z* = *−*1.67; *p* = .09).

## Discussion

In the present study, we examined whether high-definition tRNS (HD-tRNS) over the right DLPFC enhances both training outcomes and transfer to a flight simulator, while reducing workload. Contrary to prior findings [43], stimulation conferred no additional benefit over training alone for either trained task or transfer to flight simulator. No effects of stimulation were observed on learning rates for the trained tasks, nor on short-or long-term training outcomes, whether for trained-task performance, flight simulator transfer, or subjective and objective workload levels.

### tRNS does not improve direct training effects on trained tasks

Both the stimulated and the sham groups exhibited positive learning rates during the training of both MATB and SF. Likewise, both group significantly increased their global scores from baseline to the first and the last evaluation sessions in the trained tasks. Although we designed the current study to expose the participants to a substantial training period (*i.e*., 10 distributed hours of training during 5 weeks including 200 min of cumulated stimulation), no significant direct benefit of the right DLPFC stimulation has been found. At least two complementary explanations can be drawn from these observations.

First, the significant benefit of rDLPFC stimulation observed in our previous study [43] may either be marginal or a false-positive due to low sample size. However, we must specify that the protocol used in the present study was more substantial (10 hours in 5 weeks instead of 100 minutes in 5 days). Nevertheless, it is important to consider this previously observed potential benefit in practical terms. Even with a significant superiority of such a stimulation, the effect size would remain very small, thereby limiting the benefit–risk balance of stimulation for training purposes. In a recent systematic review and meta-analysis, Poppe et al. ([52]) showed that after sensitivity analyses, adding brain stimulation to cognitive training led to small, short-term improvements in learning and memory capacities compared with cognitive training alone, with no sustained long-term benefit. Although statistically significant, this effect showed substantial heterogeneity through studies. As mentioned in several recent reviews [53, 54], it is very likely that the domain of cognitive training and brain stimulation suffers from publication bias like small sample size studies, false-positive results, absence of double-blind condition and a tendency to publish only significant results [55].

A second explanation may stand in the tRNS parameters applied in the current study. We used a 100-500 Hz frequency noise with no DC-offset. The first parameter is a hardware limitation that prevents from applying a higher frequency band (up to 640Hz). It has been suggested that the propensity of the tRNS to increase cortical excitability may stand in the application of higher-frequency noise [56, 57]. It is hence possible that our 500Hz limitation reduced the tRNS effect. The second parameter could have been adopted to compare with other studies showing benefits of tRNS with a 1 or 2 mA DC-offset for instance [33, 35]. Without it, the net effect of tRNS is 0 mA (from -1 mA to 1 mA Gaussian-distribution in our case). Nevertheless, previous results showing benefits of the tRNS without DC-offset in cognitive trainings, including our own work [43] encouraged us to apply the same parameters. More recently, it has also been suggested that, in some cases, the addition of such a DC-offset may lead to a non-synergistic interaction [58] leading to counter-productive effects of the stimulation.

### tRNS does not improve transfer learning to untrained tasks

Given the absence of any stimulation effect on the trained task performance, it was unlikely that there would be a significant benefit in terms of higher transfer effect to the untrained tasks. Our results confirm this statement, since the two groups did not differ in terms of flight simulator performance. Although they both improved over sessions, no group effect has been found. This was also true for both the subjective workload assessment (NASA-TLX) and the objective assessment of auditory attentional resources (concurrent Oddball task). As discussed in a second-order meta-analysis by [14], very few studies have provided solid proof of transfer effect to untrained tasks, although they are related to the trained ones. They also showed that even when a significant transfer was shown, the effect size was very small. As a consequence, it is likely that the observed improvements between reference and short- and/or long-term sessions occurred because of time and familiarization with the simulator.

### Strengths, limits, and considerations

This study was conducted with rigorous methodology to address previous limitations. Key strengths include a double-blind procedure for both stimulation and data processing, with a true sham control group. As attested by the absence of differences between groups in the tRNS adverse effect questionnaire, we can argue that the blinding procedure was effective.

Anticipating the difficulty of recruiting private pilot license holders, we designed the study to compensate for the small sample size by increasing the number of sessions, with participants completing 10 *×* 1 hour of training and receiving a total of 200 minutes of stimulation across these sessions. The training duration, hence exceeded that of most NIBS studies in healthy populations. Finally, both direct and far transfer effects of training were assessed.

Despite careful efforts to conduct this study rigorously, several limitations should be acknowledged. First, the sample size was relatively small (30 participants, 15 per group) and mainly composed of men (26 M; 4F), resulting in limited statistical power and generalizability.

In addition, the trained-task difficulty did not change across the sessions, resulting in ceiling effects (especially visible for the MATB). This could have limited the observable benefits of brain stimulation in terms of final improvements, at least for the trained-tasks. It should however be noted that the learning rates are arguably less sensitive to this effect.

Finally, although each training session lasted one hour, brain stimulation was only applied during the first 20 minutes. Thus, any potential benefits of stimulation may have been overshadowed by the broader effects of training itself.

### Implication for pilots’ training and learning strategies

When considering the use of a stimulation tool such as transcranial brain stimulation to enhance general cognitive processes, substantial and reliable benefits would be required, particularly in the context of industrial-level training. In this study, we observed no differences between groups across sessions for either trained or untrained tasks, indicating no meaningful benefit overall.

Furthermore, applying a transcranial current to the brain—while harmless—is not without significance. The benefit-risk balance must therefore be favorable. Existing research, including applications in military operations [59] and clinical applications [60, 61], shows that the effects of the brain stimulation are either absent or typically small. These results, coupled with our findings, show a lack of consensus in the literature, and therefore call into question the practical relevance of brain stimulation tools.

## Conclusions

The current study results suggest that using HD-tRNS as a tool to enhance both cognitive training and transfer learning to more ecological multitasking tasks, such as flying a flight simulator, seems less promising when applied with the current parameters (*i.e*., 100-500Hz random noise, 4 *×* 1 HD-montage, *−*0.5 to 0.5 mA, no DC-offset).

Further research is still needed to confirm this work outcomes using different parameters and larger sample sizes. While tRNS is generally considered safe, its long-term effects and potential for misuse (e.g., cognitive enhancement outside clinical or research and training contexts) warrant careful consideration. Standardized protocols and regulatory guidelines will be essential as neuromodulation technologies become more accessible.

## Conflict of interest

The authors report no conflict of interest in the present study.

## Declaration of generative AI in scientific writing

During the preparation of this work the authors used Mistral “Le Chat” in order to rephrase some part of the manuscript and Visual Studio Copilot to generate part of the data processing code in Python. After using this tool, the authors reviewed and edited the content as needed and take full responsibility for the content of the published article.

## Funding

This work was supported by the French Government defense procurement and technology agency.

## CRediT author statement

**Sébastien Scannella:** Conceptualization, Data curation, Formal analysis, Funding acquisition, Methodology, Project administration, Resources, Software, Visualization, Supervision, Writing - Original Draft;

**Florine Riedinger:** Investigation, data collection, Methodology, Writing - Review & Editing;

**Quentin Chenot:** Conceptualization, Data curation, Formal analysis, Investigation, Methodology, Software, Supervision, Validation, Visualization, Writing - Review & Editing;

## Acknowledgements

The authors want to thank Frédéric Dehais for his help in designing the flight scenarios and Stéphane Perrey for his support in the Ethics committee approbation.

## Notes

### Competing Interest Statement

The authors have declared no competing interest.

